# Interactive Multiresolution Visualization of Cellular Network Processes

**DOI:** 10.1101/659367

**Authors:** Oscar O. Ortega, Carlos F. Lopez

**Affiliations:** Chemical and Physical Biology Program, Vanderbilt University, Nashville, Tennessee; Biochemistry Department, Vanderbilt University, Nashville, Tennessee

## Abstract

Computational models of network-driven processes have become a standard to explain cellular systems-level behavior and predict cellular responses to perturbations. Modern models can span a broad range of biochemical reactions and species that, in principle, comprise the complexity of dynamic cellular processes. Visualization plays a central role in the analysis of biochemical network processes to identify patterns that arise from model dynamics and perform model exploratory analysis. However, most existing visualization tools are limited in their capabilities to facilitate mechanism exploration of large, dynamic, and complex models. Here, we present PyViPR, a visualization tool that provides researchers static and dynamic representations of biochemical network processes within a Python-based Literate Programming environment. PyViPR embeds network visualizations on Jupyter notebooks, thus facilitating integration with Python modeling, simulation, and analysis workflows. To present the capabilities of PyViPR, we explore execution mechanisms of extrinsic apoptosis in HeLa cells. We show how community-detection algorithms can identify groups of molecular species that represent key biological regulatory functions and simplify the apoptosis network by placing those groups into interactively collapsible nodes. We then show how dynamic execution of a signal, under different kinetic parameter sets that fit the experimental data equally well, exhibit significantly different signal-execution modes in mitochondrial outer-membrane permeabilization – the point of no return in extrinsic apoptosis execution. Therefore, PyViPR aids the conceptual understanding of dynamic network processes and accelerates hypothesis generation for further testing and validation.

## INTRODUCTION

Cellular processes are controlled by networks of biomolecular interactions that process signals and trigger a response [1–3]. These molecular networks give rise to nonlinear dynamic processes that are difficult to explain and predict using reductionist methods [4]. Mathematical models of cellular signaling pathways have become commonplace in order to gain insights and describe the molecular mechanisms that control cellular processes [5–7]. In general, these models continue to grow in size and complexity, which makes the exploration of network structure and dynamics increasingly challenging. Visualization tools comprise one effective way to explore network processes and acquire conceptual insights about signal-execution mechanisms. In addition, visualization tools can facilitate detection of execution patterns, and aid in hypothesis generation for experimental validation. However, to the best of our knowledge, most tools focus on static network representations of models without strategies to deal with increasingly complex models, and generally lack support to visualize model dynamics. Therefore, there is a need for novel tools that provide viable visualizations of large models as well as support for intuitive visualizations of model dynamics.

Visualization tools can provide important insights into the complex relationships among multiple interacting components in a network. Numerous tools have been developed to generate network representations to capture relationships between model components. Some examples include molecular species networks [8], species-reactions networks [9], contact maps [10–12], model-defined rules [11, 12], and rule-based networks [13, 14], among many others [15–17]. Although these tools have been immensely useful within their domain, they exhibit limitations to visualize the structures of increasingly complex networks due to the a large number of nodes, edges, and labels. Further, standalone visualization tools can be difficult to incorporate into model building and analysis workflows, thus compounding reproducibility in analysis pipelines.

Relevant biomolecules in a signaling network process can participate in multiple dynamic interactions and exhibit nonlinear dependence on model initial conditions and kinetic parameters. Identification of reactions that drive the cellular processes is central to dynamic network analysis, yet highly challenging without visualization tools to facilitate an intuitive understanding of the signal execution mechanisms. A handful of tools to visualize dynamic network processes have been published, notably COPASI [8], and the Kappa Dynamic Influence Network (KDIN) [18]. COPASI uses a network in which nodes represent biochemical species and edges represent biochemical interactions, and it qualitatively encodes the species concentrations obtained from a simulation in the size of the box around the network nodes. Kappa employs a network in which nodes are the model rules and the edges indicate that the rules have common reactants or products species, and it quantitatively represents the temporal influence that each biochemical rule exerts on other rules. Although both tools yield useful information about dynamic network processes, information about the reactions that drive the dynamic consumption and production of different proteins is not displayed in their visualization, which is essential to understand signal execution mechanisms. Additionally, these tools have been developed for specific software environments, thus limiting their generalization for other modeling and analysis workflows.

In this work we tackle three main visualization challenges that we hope will catalyze our conceptual understanding of biological network processes: (*i*) develop legible and comprehensible visualizations of increasingly large networks; (*ii*) generate intuitive dynamic network visualizations of model simulations; and (*iii*) facilitate the integration of visualizations to model building and analysis pipelines. To tackle these challenges, we developed Python Visualization of Processes and Reactions (PyViPR), a Python framework that provides multiple static and dynamic representations of biological processes. Importantly, PyViPR unifies tools typically used in isolation, applies network community detection algorithms, and encode model simulations in node and edge properties to enable the study of large model networks and their dynamics at different resolutions. PyViPR embeds all network visualization and analysis onto Jupyter Notebooks [19] in order to facilitate reproducibility and the development of shareable model analysis pipelines. PyViPR currently supports rendering of rule-based models declared in the PySB framework [20], BioNetGen (BNG) [10] and Kappa language [11], as well as models encoded in the SBML format [21], thus providing a general tool to visualize models of biochemical network processes. In what follows we describe how PyViPR was designed and implemented, followed by a demonstration of community detection in a complex network of apoptosis execution, and finally a use case to explore dynamic execution of network processes.

## RESULTS

### Overview of PyViPR

PyViPR is a python package designed to be used within Jupyter notebooks. We chose Jupyter notebooks due to its literate programming paradigm [22] that enables the definition of both code and documentation at the same time, allowing users to develop shareable workflows for model definition, visualization, and analysis. PyViPR leverages the capabilities of PySB to generate model objects, import models from BNGL and SBML formats, and provide simulation-based results for dynamic visualization. Importantly, PyViPR brings the power of cytoscape.js [23], a well-established JavaScript library for graph visualization, to the python environment to interactively render static and dynamic visualization of model networks. In this manner, PyViPR is a software platform that integrates software packages that would traditionally be used in isolation onto a common modeling framework. Importantly, PyViPR takes advantage of community-driven software development, as it automatically accrues improvements and enhancements made to any its software components. In addition, PyViPR motivates community contributions through its open-source philosophy built around GitHub: https://github.com/LoLab-VU/PyViPR.

A typical PyViPR workflow comprises the following steps. First, a supported model file is passed to one of the PyViPR visualization functions. PyViPR then uses NetworkX [24] to convert the model components into graph nodes an edges. The user could then simplify the graph through community detection using the Louvain algorithm [25] on the NetworkX graph object. The code will then create a compound node and place all the nodes from a community within it. For dynamic visualization, PyViPR maps the simulated species concentrations and reaction data to node and edge properties. The resulting NetworkX graph is transferred to cytoscape.js via a JSON dictionary and rendered real-time in a Jupyter notebook for visualization. We note that the user can interact with all graph objects in a Jupyter notebook rendering to, e.g. change the layout, groupings, or placement of a given graph.

### Network creation from multiple model components

PyViPR supports visualization of multiple model components, including molecular species, reactions, rules, compartments, macros functions [20] and modules comprising independent model elements [20]. These components are depicted by either *simple nodes*, which are fundamental units in a graph, or *compound nodes*, that can contain children nodes and are used to group simple nodes with shared attributes or through user-defined groupings. Model species, reactions, and rules are represented by *simple nodes*, whereas model compartments, modules, and macros functions, are represented as *compound nodes*.

To build a network PyViPR obtains a list of the reactions defined in a model, adds the rules/reactions and the species involved as nodes to the network, and use edges to connect reactants and products species nodes with their respective rule/reaction node(Figure S1 A). To reduce the resolution, i.e. the number of nodes, these graphs can be projected to a unipartite graph that contains only the species or rules/reactions nodes (Figure S1 B). This unipartite species graph can then be organized by grouping the species nodes using the biological compartments on which they are located (Figure S1 C). Similarly, a unipartite rules graph can be grouped by the macro functions used to create them or the model modules where they are defined. This allows users to interactively explore and revise the model network topology at different resolutions. For a complete list of the different model components that can be visualized in a network see Figure S2.

We also added the Louvain [25] method for community detection to automatically cluster nodes and thereby simplify network complexity. Briefly, the Louvain method optimizes graph modularity by first iterating over all nodes and assigning each node to a community that results in the greatest local modularity increase, then each small community is grouped into one node and the first step is repeated until no modularity increase can occur. In this manner the Louvain algorithm finds groups of highly connected nodes that could have similar biological functions or represent molecular complex formation processes [26]. Additionally, due to the iterative nature of the Louvain algorithm, it discovers a hierarchy of communities at different scales, which can be useful for understanding the structure of a network. Finally, We embedded the detected communities into collapsible/expandable compound nodes to facilitate the exploration and navigation of large networks.

Alternatively, users can interactively define their own clusters of nodes. In this manner, the visualization becomes a dynamic process whereby users can interact in real-time with the rendered network to optimize the visualization to their needs. This so-called “human in the loop” or “active” optimization for visualization purposes could greatly accelerate how concepts are conveyed and shared by users in the systems biology community [27, 28].

### Dynamic visualization by incorporating simulation data

Models have complex dynamic patterns that arise from the temporal changes in protein concentrations and reaction rates. These dynamics are difficult to track and follow in models with large numbers of interacting proteins. Hence, the goal of PyViPR is to provide a clear and easily interpretable visualization to distinguish the interactions that drive changes in biological network processes.

PyViPR supports the dynamic visualization of results from deterministic and stochastic model simulations(Figure S1 D). This visualization mode uses a network whose nodes are the model species and the edges represent the reactions between the species. To be able to visualize the simulation in a clear and easily interpretable way we encoded the species concentrations and reaction rates into properties of nodes and edges, respectively.

To represent the temporal change in the concentration of molecular species during a simulation, we embedded pie charts inside the graph species nodes. The pie chart slices within the nodes show the concentration of a species relative to the maximum amount of the concentration attained across all time points of the simulation. The pie chart slices are updated according to the concentration of the species at each time point of a simulation. Additionally, we include information about the value of the species concentration as tooltips that can be accessed by a click-hold gesture on the species of interest.

Molecular species can be consumed or produced by the different interactions on which they are involved. Each of those interactions is represented by an edge connected to the species node. The speed at which these reactions occur changes over time and is determined by the reaction rate values obtained from a simulation. To facilitate visualization of a given molecular species consumption or production over time, each of the producing (consuming) interactions is normalized relative to the sum-total of the producing (consuming) interactions at each simulation time point. Then, the normalized values are linearly mapped to a color shades. Lighter shades mean that the flux is smaller compared to the total, conversely darker shades show that the flux is higher. Edge width represents the relative value of the reaction normalized to the maximum value that the edge can attain across all time points of the simulation. For consistency, the visualization shows how all nodes are being consumed, or alternatively, how they are produced at the same time. It is possible to change between the production and consumption visualization from a drop-down option in the rendered visualization. Finally, the simulated reaction rate values for each interaction are included as tooltips that can be accessed by a click-hold gesture on the species of interest.

Models have complex dynamic patterns that arise from the temporal changes in protein concentrations and reaction rate. These dynamics are difficult to track and follow in models with a large number of interacting proteins. Hence, the goal of this node and edge encoding is to provide a clear and easily interpretable visualization to distinguish the interactions that drive the changes in concentration of molecular species.

### PyViPR to visualize apoptosis execution

To illustrate the visualization capabilities of PyViPR, we use the Extrinsic Apoptosis Reaction Model (EARM v2.0) to study the receptor-mediated apoptosis signaling cascade.

After a ligand binds a death receptor, the apoptosis signal can flow in two ways, denoted Type I and Type II [29]. In Type I, an initiator caspase is activated which in turn activates an effector caspase. Whereas in Type II, an initiator-effector caspase cascade is initiated and induces the mitochondrial outer membrane permeabilization (MOMP) which amplifies the cell death signal. MOMP is regulated by the complex interaction of numerous pro-survival and pro-death signals. These include the Bcl-2 family of proteins, which may be anti-apoptotic (Bcl-2, Bcl-xL, Mcl1, A1, Bcl-w), pro-apoptotic (Bax, Bak, Bok), activators (Bid, Bim), and sensitizers (Bad, Puma, Noxa) [30]. The most relevant interactions from Type I and Type II apoptosis are encoded in the Extrinsic Apoptosis Reaction Model (EARM 2.0) that uses the PySB framework [20]. EARM is a relatively large model that comprises 74 molecular species, 127 parameters, 62 rules and 100 reactions.

#### Multiresolution visualization and exploration of EARM

We wanted to study the architecture of the network defined in EARM to see if it could reveal insights about the molecular organization and function in the apoptosis pathway. We started by inspecting the bipartite graph which contains 139 species and rules nodes, as well as the edges that connects the reactant species with a rule and the subsequent product species (Fig 1, EARM bipartite graph). However, this large network is difficult to explore and it does not have a discernible structure that would allow a researcher to understand how the collection of interactions between species and rules nodes drive the apoptosis mechanism. Hence, to improve model exploration we projected the species-rules bipartite graph into a species unipartite graph and grouped the nodes into communities of highly connected nodes obtained by applying the Louvain algorithm (Fig 1, Middle panel). Additionally, these communities can be interactively collapsed (Fig 1 Lower panel) to obtain a coarse-grained representation of the apoptosis pathway.

**Fig 1.**
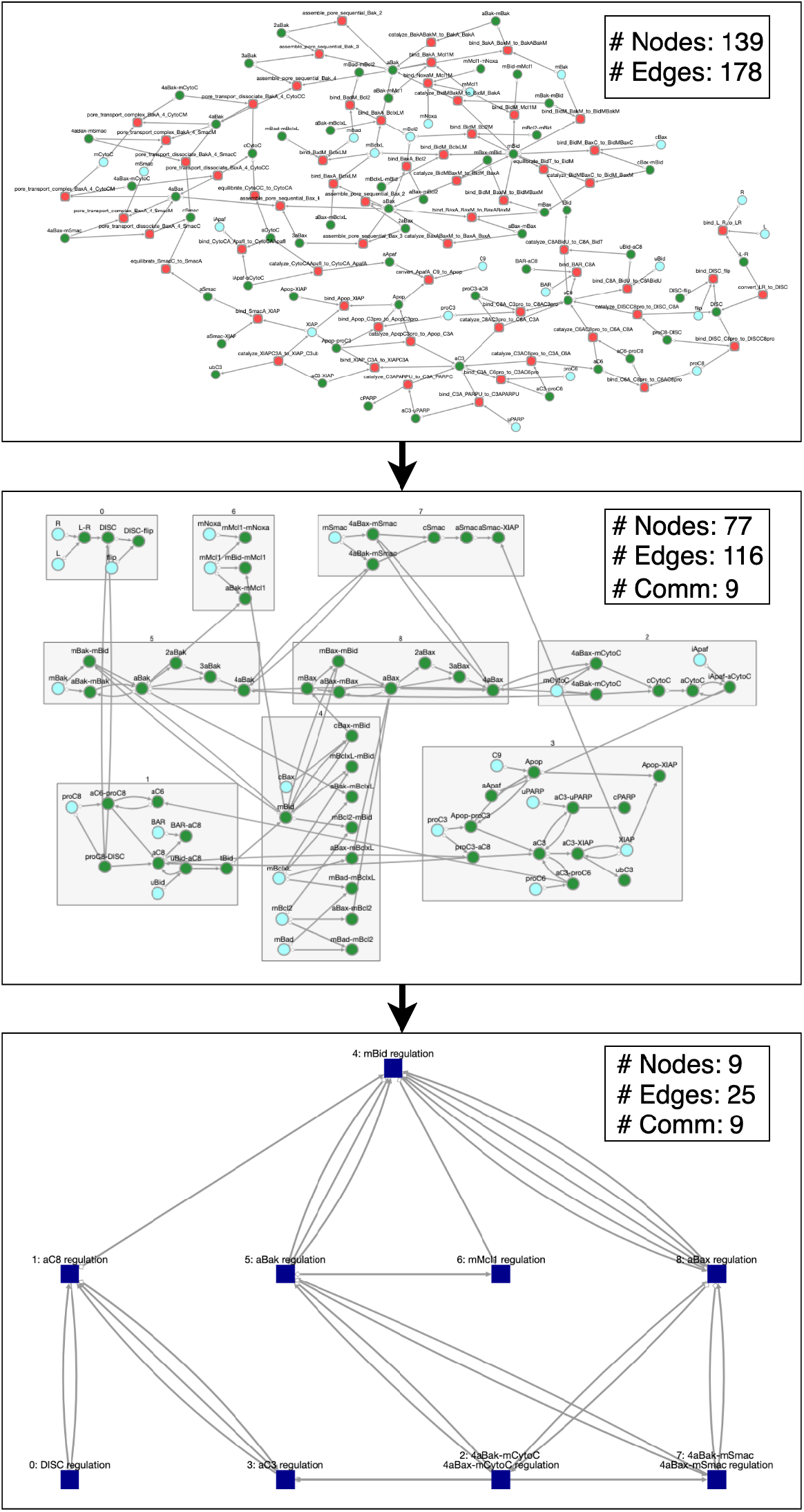
Multiscale visualization of EARM. Upper graph: EARM bipartite graph. Green nodes represent molecular species and the red nodes represent rules defined in the model. Middle graph: EARM compound graph. Densely connected nodes are grouped into communities. Lower graph: EARM communities graph. This graph is obtained by collapsing the communities into a single node. The name of the new node is determined by the species with the highest number of interactions within the community.

To visualize the species network from EARM and the communities detected by the Louvain algorithm we used the PyViPR function *sp_comm_view*. The community detection algorithm found 9 communities, labeled 0-8 (Figure S3). Community 0 contains the ligand-receptor interactions that lead to the DISC formation and regulation by Flip [31]. It is uniquely connected to community 1 that comprises initiator Caspase 8 (C8) activation by DISC and Caspase 6 and the subsequent truncation of Bid by activated C8 [32]. Community 1 is particularly interesting because it is connected to communities 3 and 4 which are the starting points for differentiation between Type I and Type II apoptosis, respectively. This suggests that the proteins in Community 1 play an important role in apoptotic signaling as their interactions could determine the type of apoptosis executed by a cell [33].

Community 4 includes the regulation of mitochondrial Bid (mBid), Bax and Bak by the antiapoptotic proteins BclxL and Bcl2 [34], and it is connected to Community 6 that corresponds to the interactions of the anti-apoptotic protein Mcl1. Additionally, Community 4 is also connected to communities 5 and 8 that correspond to Bak and Bax activation, polymerization and pore formation, respectively. The interactions among these communities describes the overall regulation at the mitochondria that lead to MOMP formation, the point-of-no-return in Type II apoptosis execution. The MOMP-related communites are connected to communities 2 and 7, which correspond to the release of Cytochrome c and Smac from the MOM through pores made by Bax and Bak. Finally, these communities connect to Community 3 which corresponds to the activation of executioner Caspase 3 (C3). As shown, C3 can be directly activated by C8 (Type I) or by the apoptosome that is formed after Cytochrome c is released from the MOM (Type II).

We found that the Louvain community detection algorithm grouped biochemical species into biologically relevant functional processes of key protein activation and regulation during the apoptosis signaling pathway. This suggests that community detection could be used as a dynamic “coarse-graining” methodology to automatically group biochemical interactions in a network and simplify mechanism exploration. The community detection analysis showed that C8 and C3 are the species with highest node degree in their respective communities, indicating that they are likely essential for the regulation of biochemical events happening in their communities. We also found that mBid has the highest node degree of interactions with other communities indicating that it is likely relevant to transmit information across pathway segments.

#### Parameter sets fit experimental data but yield different network dynamics in EARM

Systems biology models are sloppy [35], which means that only changes in a few parameter combinations significantly affect model output while other parameters may vary over a wide range of values without significantly affecting model output. As a result, when models are calibrated there are many different parameter sets that fit the experimental data equally well [36]. Therefore, we decided to investigate the mechanistic implications of these different fitted parameters sets on apoptosis signal execution. We focused on mitochondrial Bid as its dynamics is tightly linked to MOMP, making it one of the most important regulators of time-to-death of cells [37]

We calibrated EARM to previously published experimental data [37] using a Particle Swarm Optimization algorithm [38, 39], and obtained 6572 different parameter sets that fit equally well with a sum of squares error equal or less than 2.8. We applied the z-score to standardize the parameter sets and then calculated all pairwise dissimilarities using the euclidean distance. Next, we chose the two maximally different parameter sets (Table S4), labelled parameter set 1 and parameter set 2, to study their effect in the mBid interactions dynamics (Fig 2 EARM calibrated to experimental data). These dynamics are dictated by the interactions of Bid with various anti-apoptotic and pro-apoptotic proteins [30], which are difficult to simultaneously track and follow over time.

**Fig 2.**
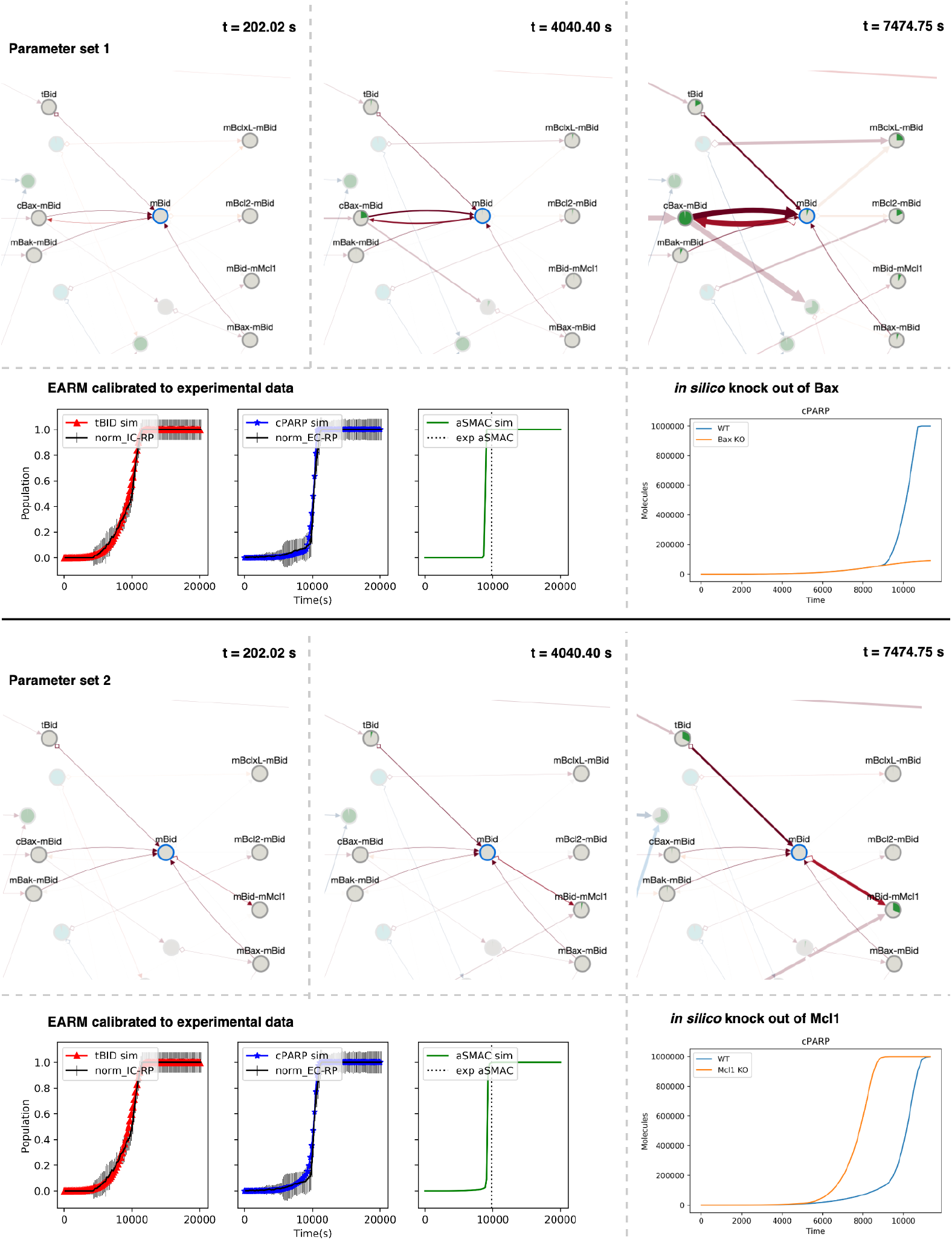
Dynamic visualization and analysis of the interactions of mitochondrial Bid with two calibrated parameter sets. The upper panel includes three snapshots that show the temporal changes in strength of the interactions between mitochondrial Bid and the anti-apoptotic and pro-apoptotic proteins. The lower left panel shows how the model simulations with the calibrated parameter set reproduces the time-course data from the experiments. Finally, the lower right panel corresponds to a *in silico* knock out of the protein involved in the dominant reaction rate observed from the dynamic visualization.

We utilized the PyViPR *sp_dyn_view* function to visualize how the signal initialized by the TNF-related apoptosis-inducing ligand (TRAIL) propagated through the EARM network and focused on flow through the MOMP module. This visualization allowed us to identify the molecular reactions that most rapidly consume mBid, indicating that they are potential targets to effectively modulate the time-to-death of cells. For parameter set 1, we observed that most of mBid was used to transport cytosolic Bax to the MOM while no activation of Bak occurred, indicating that the pores in the MOM were primarily made by Bax and that the model with this parameter set is particularly sensitive to Bax inhibition. In contrast, for parameter set 2 we observed that mBid activity was inhibited primarily by the anti-apoptotic protein Mcl1, indicating that it plays an important role in throttling MOMP (Fig 2 A and B, upper panel). To verify our visualization-based analysis, we carried out two in-silico experiments. First, we did a Bax knockout and ran a simulation of EARM with parameter set 1. We found that knocking out Bax protected cells from apoptosis induction with TRAIL, confirming that Bax has an essential role in apoptosis. Second, we did an Mcl1 knockout and ran a simulation of EARM with parameter set 2. We found that the time-to-death was reduced by 22.6%, corroborating that Mcl1 was delaying the apoptosis execution by binding to mBid. These results demonstrated that although these two parameter sets fit the data equally well, they executed the apoptosis signal in different ways; specifically, in this case, the parameter sets determined whether Bax or Mcl1 played the key role in regulating apoptosis execution. These observations align we experimental results that showed how cell lines might depend on different proteins for apoptosis execution [40, 41]. Therefore, visualization of the dynamic processes enabled us to identify key reactions under different parameter sets and generate testable hypotheses used to further understand apoptosis execution mechanisms. This same paradigm could be used to generate model-driven hypothesis to test experimentally.

## DISCUSSION

In this paper we present PyViPR, a novel tool to visualize the structure and dynamics of biochemical models. PyViPR integrates a community detection algorithm to organize the nodes of biochemical networks to improve the legibility of large networks. PyViPR provides an intuitive dynamic visualization that facilitates the identification of dominant reaction rates that are either consuming or producing proteins.

To illustrate the capabilities of PyViPR, we provided empirical evidence of how visualization of biochemical reaction networks at multiple resolutions and with model dynamics could quickly lead to biological knowledge and hypothesis generation. Visualization of EARM communities (Fig 1) enabled us to identify groups of molecular species that are functionally related to important processes during apoptosis execution, and to the formation of protein complexes. This network organization by communities can show how a specific outcome can be attained through different processes (communities), thus providing an efficient way to identify potential targets to control the signal in complex biochemical networks. Furthermore, Visualization of EARM dynamics enabled us to identify key regulatory proteins in the apoptosis execution process and their dependence in specific parameter values (Fig 2). Overall, PyViPR enables researchers to explore hypotheses about the dynamic regulation of biochemical models within an interactive setting that opens the door for novel workflow paradigms.

We believe that PyViPR could be incorporated onto existing modeling and simulation workflows such as those provided by Python-based tools such as Tellurium notebooks [42] and PySCeS [43]. In addition, non-Python users could also take advantage of PyViPR using SBML import, thus providing a generalizable tool for model exploration. In the future, we plan to incorporate more complex community-detection algorithm that consider the weight of the edges for the clustering of nodes. Additionally, we plan to improve the synchronization from the JavaScript frontend to the Python backend to enable users to interactively modify model parameters and components.

All the model exploratory analyses, which includes model calibration, visualization, hypothesis exploration, and testing, were performed in Jupyter Notebooks. These notebooks, which are shareable and reusable, contain all the source code, and markup text that explains the rationale for each step in the analysis (See Supplement Section and LINK). This promotes reproducibility and transparency by enabling other researchers to rerun or expand the presented model analysis. We invite the community to contribute to open-source tools such as PyViPR to improve model analysis and visualization

## Supporting information

Table S4 parameter set 1

Table S4 parameter set 2

## STAR⋆METHODS

### KEY RESOURCES TABLE

| REAGENT or RESOURCE | SOURCE | IDENTIFIER |
| --- | --- | --- |
| Deposited Data |  |  |
| HeLa cells FRET reporter proteins data | [37] | https://github.com/sorgerlab/earm |
| Software and Algorithms |  |  |
| PyViPR | This paper | https://github.com/LoLab-VU/pyvipr |
| PySB | [20] | http://pysb.org/ |
| Networkx | [24] | https://networkx.github.io |
| Louvain algorithm | [25] | https://github.com/taynaud/python-louvain |
| cytoscape.js | [23] | http://js.cytoscape.org/ |

### CONTACT FOR REAGENT AND RESOURCE SHARING

Further information and requests for resources and reagents should be directed to and will be fulfilled by the Lead Contact, Carlos F. Lopez

### METHOD DETAILS

PyViPR was developed using Python and JavaScript. The Python package PySB [20] was used to generate biochemical models, and import Bionetgen and SBML models. Networkx [24] was used to generate node edge graphs that store information about model components and simulation results. The louvain algorithm [25] was used to cluster densely connected node in networks.

The JavaScript package cytoscape.js [23] was used to render NetworkX graphs, apply layout algorithms to the rendered networks, and enable the dynamic visualization of model simulation results. JavaScript was used to embed the resulting network into a Jupyter Notebook [19].

## DATA AND SOFTWARE AVAILABILITY

The python package PyViPR is an open-source project under the MIT License. Stable releases of PyViPR are available on pypi and the latest unreleased version can be downloaded from github https://github.com/LoLab-VU/PyViPR. The documentation with examples and description of the available functions is available at https://PyViPR.readthedocs.io. A Jupyter notebook with the code to reproduce all the figures included in the manuscript can be found in binder https://mybinder.org/v2/gh/LoLab-VU/PyViPR/master.

### Supporting information

**Figure S1.**
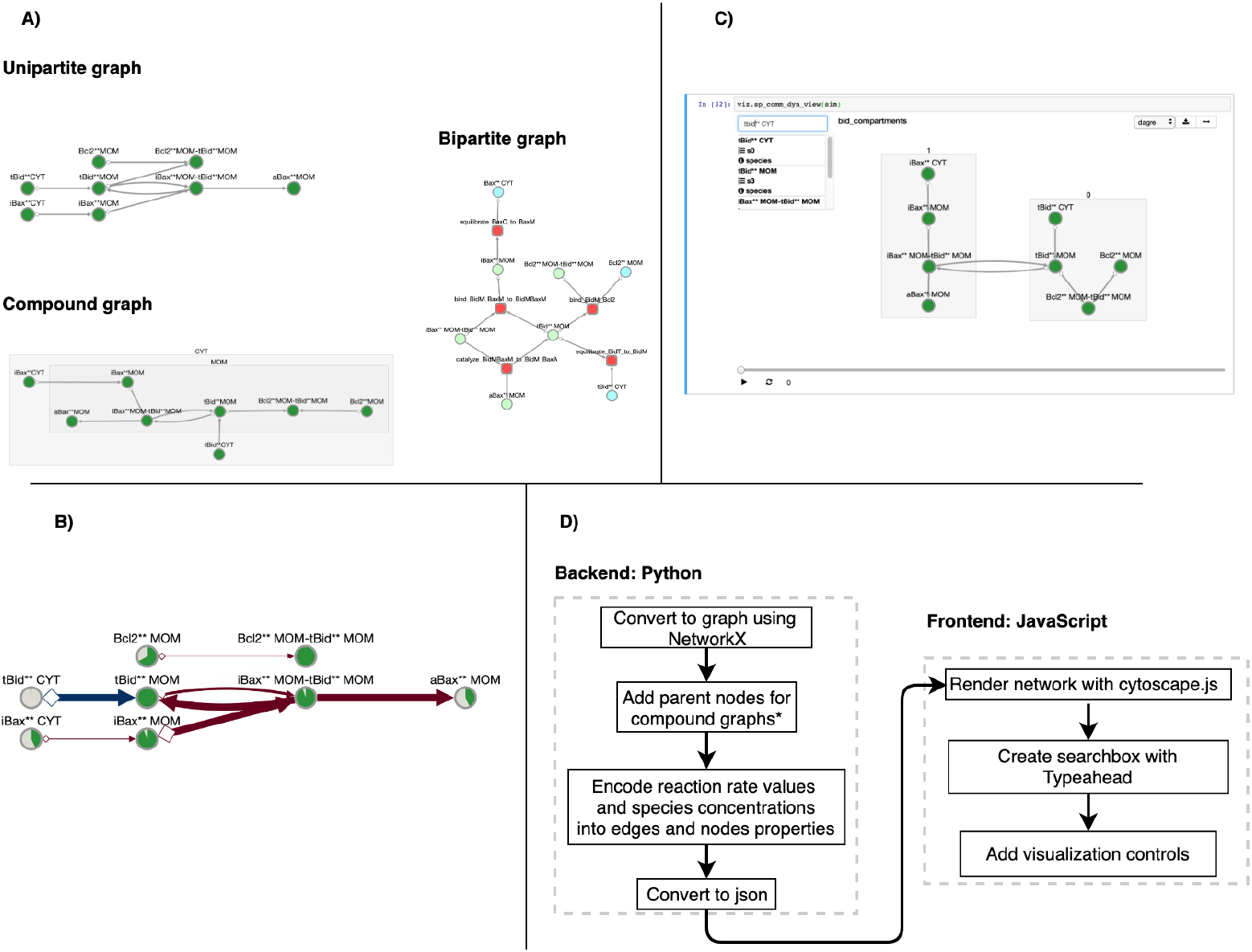
Network visualization modes in PyViPR. PyViPR supports four major modes of network visualization. (A) A bipartite graph where one set of nodes represents the model species, the second set of nodes represents model model rules, and the edges connect reactant and product species with their corresponding rule. (B) A unipartite graph where each node represents a chemical species and edges represent biochemical interactions. (C) A compound graph where the nodes are grouped by the compartments on which they are located. (D) Snapshot of dynamic visualization in a unipartite graph. Nodes represent chemical species, edges represent biochemical reactions, and the pie charts inside nodes represent species concentration over time.

**Figure S2.**
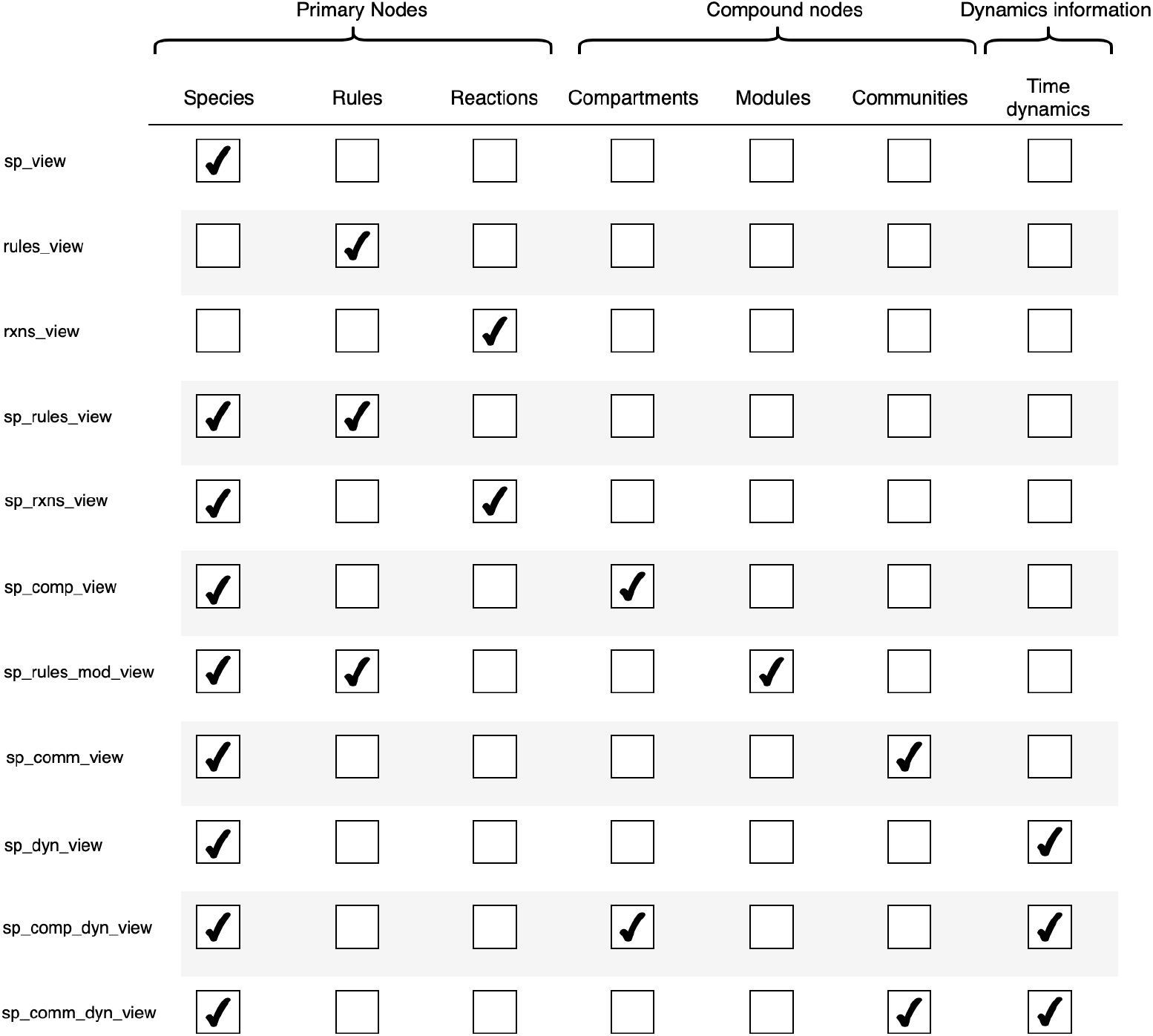
Names of functions to create model visualizations and the model components included in a network. There are three types of model components that can be used for visualization purposes. The first one are the primary nodes which correspond to model species, rules and reactions. Second is the compound nodes that include model compartment, modules/files, and communities detected by clustering algorithms. Finally, we have the dynamics information which correspond to the simulation results of a model. The marked check-boxes for each of the functions are the information displayed in a network.

**Figure S3.**
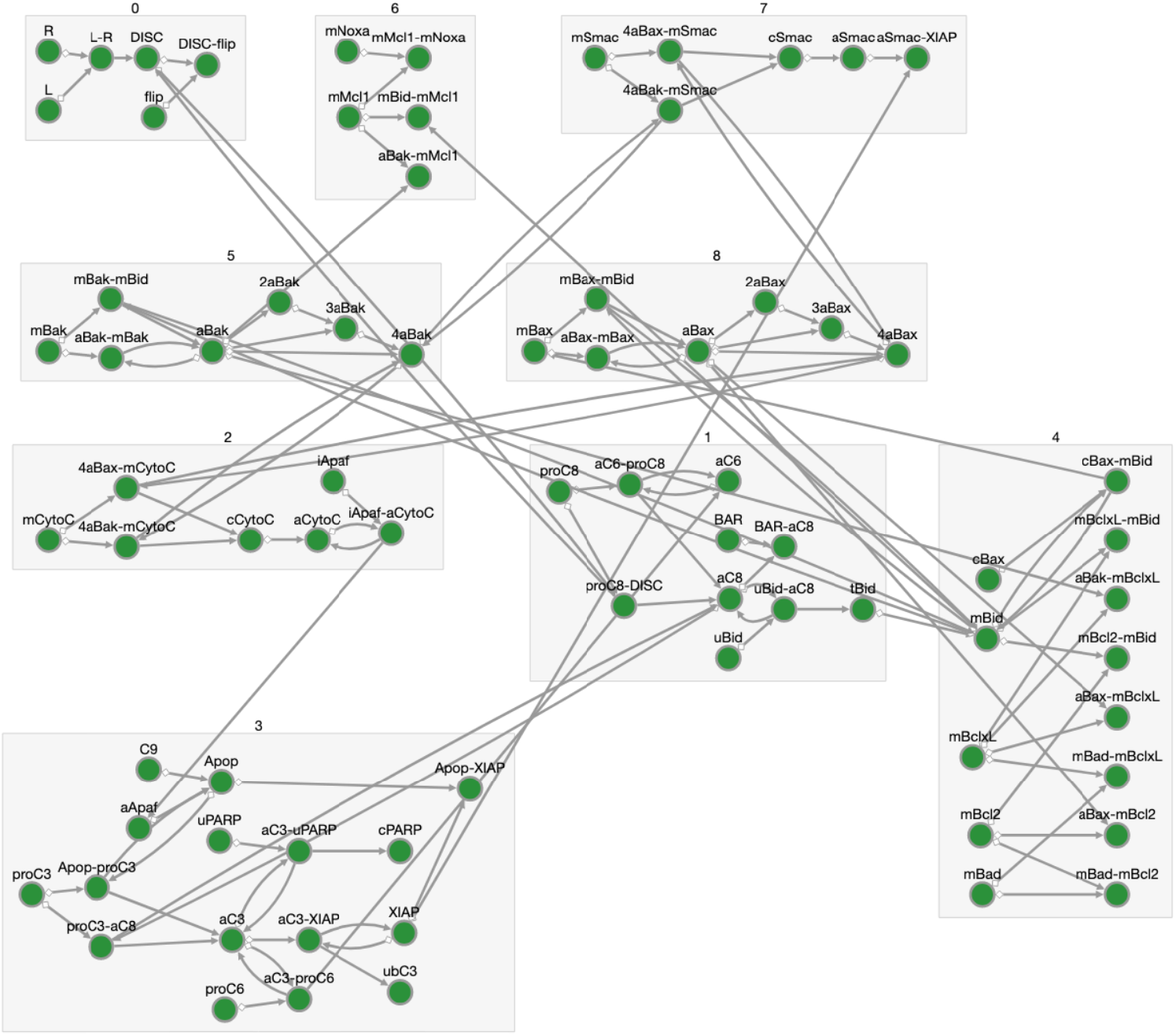
Communities detected in EARM. Species network of EARM. Each of the nodes represents a molecular species defined in the model, and the edges depict the interactions between species. Species nodes are clustered in 8 groups. These are the communities, labeled from 0 to 7, detected by the Louvain algorithm

**Table S4. Values of EARM calibrated parameters**

**Video S5. Video of signal execution for parameter set 1 in EARM.**

**Video S6. Video of signal execution for parameter set 2 in EARM.**

## Acknowledgments

We thank Blake Wilson, Leonard Harris, and Alexander Lubbock for their useful insights throughout the development of PyViPR and for useful feedback in writing this manuscript. This work was supported by NSF award MCB 1411482 to CFL, NIH-U01 award 1U01CA215845 to CFL, the Vanderbilt International Students Program to OOO, and the Vanderbilt Pearson Graduate Fellowship to OOO.

## References

1. Lemmon MA, Schlessinger J. Cell signaling by receptor tyrosine kinases. Cell. 2010;141(7):1117–34. doi:10.1016/j.cell.2010.06.011.

2. Blinov ML, Faeder JR, Goldstein B, Hlavacek WS. A network model of early events in epidermal growth factor receptor signaling that accounts for combinatorial complexity. BioSystems. 2006;83(2-3 SPEC. ISS.):136–151. doi:10.1016/j.biosystems.2005.06.014.

3. Sachs K, Perez O, Pe’er D, Lauffenburger DA, Nolan GP. Causal protein-signaling networks derived from multiparameter single-cell data.[see comment][erratum appears in Science. 2005 Aug 19;309(5738):1187]. Science. 2005;308(5721):523–529.

4. Ahn AC, Tewari M, Poon CS, Phillips RS. The Limits of Reductionism in Medicine: Could Systems Biology Offer an Alternative? PLoS Medicine. 2006;3(6):e208. doi:10.1371/journal.pmed.0030208.

5. Gaddy TD, Wu Q, Arnheim AD, Finley SD. Mechanistic modeling quantifies the influence of tumor growth kinetics on the response to anti-angiogenic treatment. PLoS Computational Biology. 2017;13(12):1–23. doi:10.1371/journal.pcbi.1005874.

6. Perry NA, Kaoud TS, Ortega OO, Kaya AI, Marcus DJ, Pleinis JM, et al. Arrestin-3 scaffolding of the JNK3 cascade suggests a mechanism for signal amplification. Proceedings of the National Academy of Sciences. 2019;116(3):810–815. doi:10.1073/pnas.1819230116.

7. Albeck JG, Burke JM, Spencer SL, Lauffenburger DA, Sorger PK. Modeling a Snap-Action, Variable-Delay Switch Controlling Extrinsic Cell Death. PLoS Biology. 2008;6(12):e299. doi:10.1371/journal.pbio.0060299.

8. Bergmann FT, Hoops S, Klahn B, Kummer U, Mendes P, Pahle J, et al. COPASI and its applications in biotechnology. Journal of Biotechnology. 2017;261(June):215–220. doi:10.1016/j.jbiotec.2017.06.1200.

9. Schaff JC, Vasilescu D, Moraru II, Loew LM, Blinov ML. Rule-based modeling with Virtual Cell. Bioinformatics. 2016;32(18):2880–2882. doi:10.1093/bioinformatics/btw353.

10. Harris LA, Hogg JS, Tapia JJ, Sekar JAP, Gupta S, Korsunsky I, et al. BioNetGen 2.2: Advances in rule-based modeling. Bioinformatics. 2016;32(21):3366–3368. doi:10.1093/bioinformatics/btw469.

11. Boutillier P, Maasha M, Li X, Medina-Abarca HF, Krivine J, Feret J, et al. The Kappa platform for rule-based modeling. Bioinformatics. 2018;34(13):i583–i592. doi:10.1093/bioinformatics/bty272.

12. Cheng HC, Angermann BR, Zhang F, Meier-Schellersheim M. NetworkViewer: Visualizing biochemical reaction networks with embedded rendering of molecular interaction rules. BMC Systems Biology. 2014;8(1):1–16. doi:10.1186/1752-0509-8-70.

13. Smith AM, Xu W, Sun Y, Faeder JR, Marai GE. RuleBender: integrated modeling, simulation and visualization for rule-based intracellular biochemistry. BMC Bioinformatics. 2012;13(Suppl 8):S3. doi:10.1186/1471-2105-13-S8-S3.

14. Danos V, Feret J, Fontana W, Harmer R, Hayman J, Krivine J, et al. Graphs, Rewriting and Pathway Reconstruction for Rule-Based Models. FSTTCS 2012 – IARCS Annual Conference on Foundations of Software Technology and Theoretical Computer Science. 2012;18(Fsttcs):276–288. doi:10.4230/LIPIcs.FSTTCS.2012.276.

15. Kolpakov FA, Puzanov MV, Koshukov A, Ras S. BioUML: Visual modeling, automated code generation and simulation of biological systems; 2006.

16. Tiger CF, Krause F, Cedersund G, Palmér R, Klipp E, Hohmann S, et al. A framework for mapping, visualisation and automatic model creation of signal-transduction networks. Molecular Systems Biology. 2012;8(578):1–20. doi:10.1038/msb.2012.12.

17. Dang TN, Murray P, Aurisano J, Forbes AG. ReactionFlow: an interactive visualization tool for causality analysis in biological pathways. BMC Proceedings. 2015;9(6):S6. doi:10.1186/1753-6561-9-S6-S6.

18. Forbes AG, Burks A, Lee K, Li X, Boutillier P, Krivine J, et al. Dynamic Influence Networks for Rule-based Models. IEEE Transactions on Visualization and Computer Graphics. 2017;2626(c). doi:10.1109/TVCG.2017.2745280.

19. Kluyver T, Ragan-Kelley B, Pérez F, Granger BE, Bussonnier M, Frederic J, et al. Jupyter Notebooks – a publishing format for reproducible computational workflows. In: ELPUB; 2016.

20. Lopez CF, Muhlich JL, Bachman JA, Sorger PK. Programming biological models in Python using PySB. Molecular Systems Biology. 2013;9(646):1–19. doi:10.1038/msb.2013.1.

21. Hucka M, Finney A, Sauro HM, Bolouri H, Doyle JC, Kitano H, et al. The systems biology markup language (SBML): A medium for representation and exchange of biochemical network models. Bioinformatics. 2003;19(4):524–531. doi:10.1093/bioinformatics/btg015.

22. Knuth DE. Literate Programming. The computer journal. 2001;27(2):97–111.

23. Franz M, Lopes CT, Huck G, Dong Y, Sumer O, Bader GD. Cytoscape.js: A graph theory library for visualisation and analysis. Bioinformatics. 2015;32(2):309–311. doi:10.1093/bioinformatics/btv557.

24. Hagberg AA, Schult DA, Swart PJ. Exploring network structure, dynamics, and function using NetworkX. Proceedings of the 7th Python in Science Conference (SciPy). 2008;(SciPy):11–15. doi:10.1016/j.jelectrocard.2010.09.003.

25. Blondel VD, Guillaume JL, Lambiotte R, Lefebvre E. Fast unfolding of communities in large networks. Journal of Statistical Mechanics: Theory and Experiment. 2008;2008(10). doi:10.1088/1742-5468/2008/10/P10008.

26. Fortunato S. Community detection in graphs. Physics Reports. 2010;486(3-5):75–174. doi:10.1016/j.physrep.2009.11.002.

27. Daschinger M, Knote A, Green R, Von Mammen S. A human-in-the-loop environment for developmental biology. The 2018 Conference on Artificial Life: A Hybrid of the European Conference on Artificial Life (ECAL) and the International Conference on the Synthesis and Simulation of Living Systems (ALIFE). 2017;(29):475–482. doi:10.1162/isala078.

28. Holzinger A. Interactive machine learning for health informatics: when do we need the human-in-the-loop? Brain Informatics. 2016;3(2):119–131. doi:10.1007/s40708-016-0042-6.

29. ö zören N, El-Deiry WS. Defining characteristics of types I and II apoptotic cells in response to TRAIL. Neoplasia. 2002;4(6):551–557. doi:10.1038/sj.neo.7900270.

30. Czabotar PE, Lessene G, Strasser A, Adams JM. Control of apoptosis by the BCL-2 protein family: implications for physiology and therapy. Nature reviews Molecular cell biology. 2014;15(1):49–63. doi:10.1038/nrm3722.

31. Safa AR. c-FLIP, A MASTER ANTI-APOPTOTIC REGULATOR HHS Public Access. Exp Oncol. 2012;34(3):176–184.

32. Li H, Zhu H, Xu CJ, Yuan J. Cleavage of BID by caspase 8 mediates the mitochondrial damage in the Fas pathway of apoptosis. Cell. 1998;94(4):491–501. doi:10.1016/S0092-8674(00)81590-1.

33. Shelton SN, Dillard CD, Robertson JD. Activation of caspase-9, but not caspase-2 or caspase-8, is essential for heat-induced apoptosis in jurkat cells. Journal of Biological Chemistry. 2010;285(52):40525–40533. doi:10.1074/jbc.M110.167635.

34. Kale J, Osterlund EJ, Andrews DW. BCL-2 family proteins: changing partners in the dance towards death. Cell Death and Differentiation. 2017;25(1):65–80. doi:10.1038/cdd.2017.186.

35. Gutenkunst RN, Waterfall JJ, Casey FP, Brown KS, Myers CR, Sethna JP. Universally sloppy parameter sensitivities in systems biology models. PLoS Computational Biology. 2007;3(10):1871–1878. doi:10.1371/journal.pcbi.0030189.

36. Eydgahi H, Chen WW, Muhlich JL, Vitkup D, Tsitsiklis JN, Sorger PK. Properties of cell death models calibrated and compared using Bayesian approaches. Molecular systems biology. 2013;9(644):644. doi:10.1038/msb.2012.69.

37. Spencer SL, Gaudet S, Albeck JG, Burke JM, Sorger PK. Non-genetic origins of cell-to-cell variability in TRAIL-induced apoptosis. Nature. 2009;459(7245):428–432. doi:10.1038/nature08012.

38. Kennedy J, Eberhart R. Particle swarm optimization. Neural Networks, 1995 Proceedings, IEEE International Conference on. 1995;4:1942–1948 vol.4. doi:10.1109/ICNN.1995.488968.

39. Pino J. LoLab-VU/ParticleSwarmOptimization: simplePSO release 1.0; 2019. Available from: https://doi.org/10.5281/zenodo.2612913.

40. Deng Y, Lin Y, Wu X. TRAIL-induced apoptosis requires Bax-dependent mitochondrial release of Smac/DIABLO. Genes and Development. 2002;16(1):33–45. doi:10.1101/gad.949602.

41. Zhou P, Qian L, Kozopas KM, Craig RW. Mcl-1, a Bcl-2 family member, delays the death of hematopoietic cells under a variety of apoptosis-inducing conditions. Blood. 1997;89(2):630–43.

42. Medley JK, Choi K, KÃ¶nig M, Smith L, Gu S, Hellerstein J, et al. Tellurium notebooks-An environment for reproducible dynamical modeling in systems biology. PLOS Computational Biology. 2018;14(6):1–24. doi:10.1371/journal.pcbi.1006220.

43. Olivier BG, Rohwer JM, Hofmeyr JHS. Modelling cellular systems with PySCeS. Bioinformatics. 2004;21(4):560–561. doi:10.1093/bioinformatics/bti046.

